# Glucocerebrosidase deficiency leads to neuropathology via cellular immune activation

**DOI:** 10.1101/2023.12.13.571406

**Authors:** Evelyn S. Vincow, Ruth E. Thomas, Gillian Milstein, Gautam Pareek, Theo Bammler, James MacDonald, Leo Pallanck

## Abstract

Mutations in *GBA* (*glucosylceramidase beta*), which encodes the lysosomal enzyme glucocerebrosidase (GCase), are the strongest genetic risk factor for the neurodegenerative disorders Parkinson’s disease (PD) and Lewy body dementia. Recent work has suggested that neuroinflammation may be an important factor in the risk conferred by *GBA* mutations. We therefore systematically tested the contributions of immune-related genes to neuropathology in a *Drosophila* model of GCase deficiency. We identified target immune factors via RNA-Seq and proteomics on heads from GCase-deficient flies, which revealed both increased abundance of humoral factors and increased macrophage activation. We then manipulated the identified immune factors and measured their effect on head protein aggregates, a hallmark of neurodegenerative disease. Genetic ablation of humoral (secreted) immune factors did not suppress the development of protein aggregation. By contrast, re-expressing *Gba1b* in activated macrophages suppressed head protein aggregation in *Gba1b* mutants and rescued their lifespan and behavioral deficits. Moreover, reducing the GCase substrate glucosylceramide in activated macrophages also ameliorated *Gba1b* mutant phenotypes. Taken together, our findings show that glucosylceramide accumulation due to GCase deficiency leads to macrophage activation, which in turn promotes the development of neuropathology.

**Author Summary:** Mutations in the gene *GBA* are the largest risk factor for developing Parkinson’s disease and Lewy body dementia, diseases in which important brain cells die. We know that the immune system can be involved in these diseases, and that *GBA* mutations cause immune changes. We did experiments to learn how the immune system changes could make brain cells more likely to die. Using a fruit fly that was missing the fly version of *GBA*, we found out that inappropriately activated immune cells, but not secreted immune proteins, were important in the development of brain problems. We also learned that the abnormal activation was triggered by the lack of *GBA* function in the immune cells, not by signals from the brain or other parts of the body. We would like to find out next whether the immune cells get inside the brain or cause harm from a distance. What we learned matters because it could help us prevent or cure brain diseases associated with *GBA* mutations. Treating the abnormal activation of immune cells in people with these mutations might help prevent damage to the brain.

## Introduction

The single strongest genetic risk factor for both Parkinson’s disease (PD) and the related disorder Lewy body dementia is mutations in *GBA* [1–3], which encodes the lysosomal enzyme glucocerebrosidase (glucosylceramidase; GCase). Heterozygous *GBA* mutations increase PD risk, while homozygous mutations cause Gaucher disease (GD) [1, 2], which is marked by varying degrees of neuropathology [4]. Exactly how *GBA* mutations increase the risk of neuropathology and neurodegeneration is not clear. In recent years, however, increasing evidence has indicated a connection between *GBA* mutations and immune system abnormalities [5–7]. Neuroinflammation has been implicated more generally in the pathogenesis of Parkinson’s disease [8, 9], Lewy body dementia [10], and other neurodegenerative disorders [11]. Studies of GCase-deficient humans and animals have revealed increases in cytokine and chemokine abundance, both within and outside the central nervous system [12]. Also, abnormalities of peripheral immune cells such as macrophages are a hallmark of GD, including the classic morphological defects attributed to accumulation of glucosylceramide (GlcCer), the substrate of GCase [13, 14]. Recent reports analyzing peripheral immune cells from humans with *GBA* mutations have described multiple defects including excessive activation, impaired phagocytosis, and hyperreactivity to pathogen-associated molecular patterns [15–18]. Both homozygous and heterozygous mutations in *GBA* have been shown to cause large-scale transcriptional alterations, including increases in proinflammatory transcripts [7]. These transcriptomic studies provide a rich source of data, but do not indicate whether the immune abnormalities are pathogenic.

To investigate whether immune alterations are involved in the development of neurological phenotypes in GCase-deficient organisms, we used our previously created *Drosophila* model of *GBA* deficiency, the *Gba1b* mutant [19]. This mutant recapitulated key neurodegenerative phenotypes including the accumulation of insoluble ubiquitinated proteins and impaired locomotion [19, 20]. The major humoral immune pathways in mammals, including Toll, NF-κB/Imd, and JAK/STAT, are conserved in *Drosophila* [21, 22], and *Drosophila* blood cells (hemocytes) are a well-validated model for macrophages and other human myeloid immune cells [23, 24]. In addition, *Drosophila* studies have shown the same association between GCase deficiency and immune system activation that is seen in mammals [25, 26]. We therefore used the *Gba1b* mutant to determine which components of the immune system contribute to the neuropathology associated with GCase deficiency.

We began our investigation with RNA-Seq on heads from *Gba1b* mutants and controls. We found strong increases in immune-related transcripts, including increased abundance of humoral immune factors and activated macrophage markers. When we genetically suppressed the expression of humoral immune factors in *Gba1b* mutants, there was no decrease in the accumulation of insoluble ubiquitinated protein. Manipulation of cellular immunity, by contrast, did decrease the development of *Gba1b* mutant pathology. Restoring *Gba1b* expression or reducing GlcCer production in activated macrophages ameliorated the mutants’ biochemical and behavioral phenotypes. Together, our findings implicate the accumulation of GlcCer in the activation of peripheral immune cells, which in turn promotes the neurodegeneration associated with GCase deficiency.

## Results

### *Gba1b* mutants have increased abundance of innate immune factors

To identify immune-related genes that might contribute to *Gba1b* mutant phenotypes, we performed RNA-Seq on samples from the heads of *Gba1b* mutants and controls. We detected 10,903 transcripts, of which 379 (3.5%) were significantly increased in abundance in *Gba1b* mutants, while 175 (1.6%) were decreased in abundance. The transcripts most strikingly increased in abundance in *Gba1b* mutants included multiple members of the *Turandot* family, a group of genes transcribed in response to immune threats and stressors [27, 28], and the *Tep* (thioester-containing protein) family (Fig 1A), which is orthologous to the mammalian complement system [29, 30]. Enrichment analysis with PANGEA [31] revealed that the transcripts increased in abundance in *Gba1b* mutants were strongly enriched for immune- and defense-related functions (Fig 1B-C).

**Figure 1.**
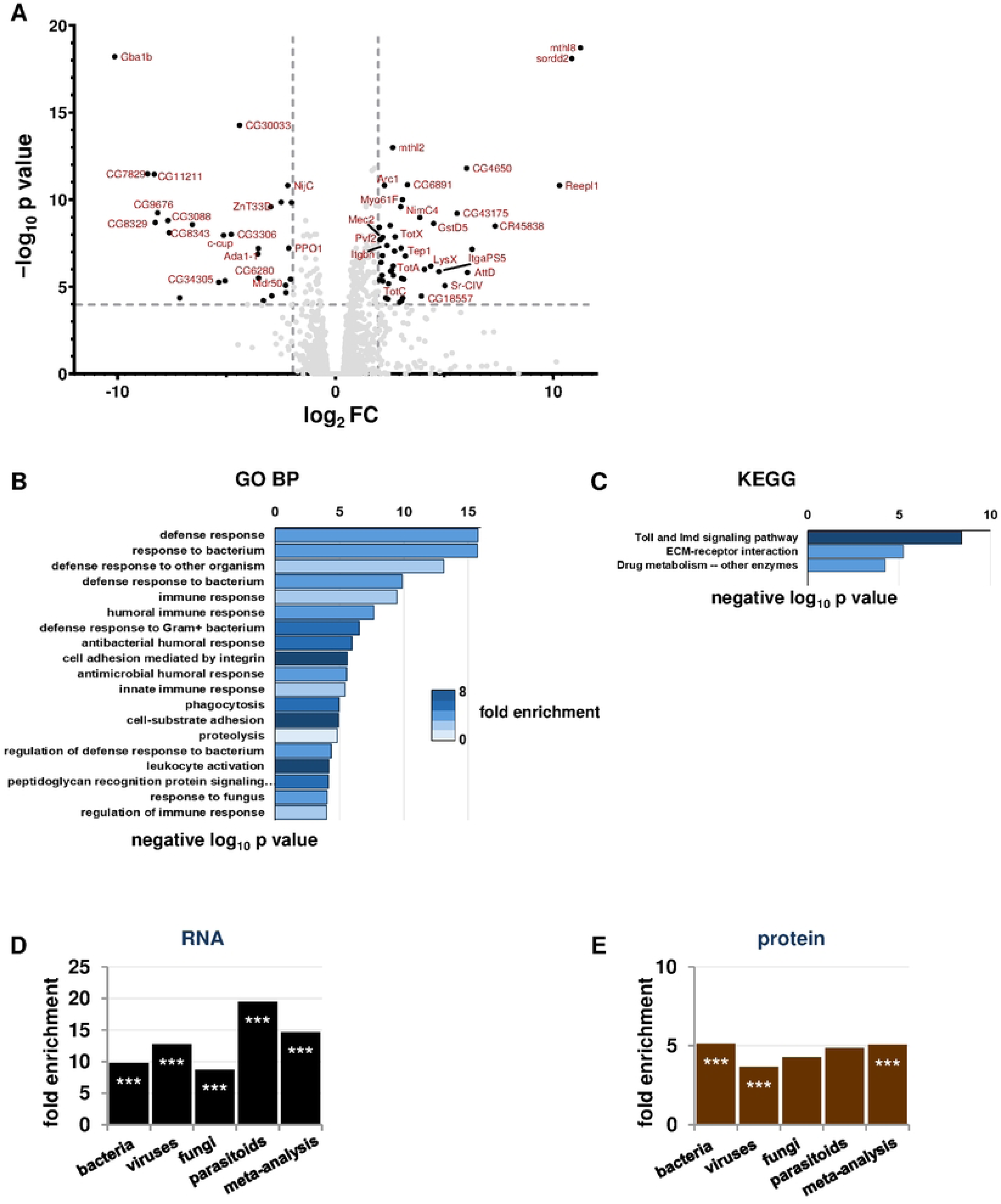
RNA-Seq and proteomics demonstrate generalized immune activation in *Gba1b* mutants. (A) Volcano plot of RNA-Seq fold change data in heads from *Gba1b* mutants vs. controls. Dashed lines mark log_2_FC of 2 and negative log_10_ *p* value of 4. (B-C) Plots of GO Biological Process (B) and KEGG Pathways terms (C) from PANGEA enrichment analysis of *Gba1b* RNA-Seq data. (D-E) Comparison of transcripts (D) and proteins (E) with increased abundance in *Gba1b* mutants to transcripts with increased abundance in immune challenge studies (see Materials and Methods for details). \*\*\**p* < 0.005 by Fisher exact test, indicating a significantly greater correspondence between *Gba1b* mutant transcriptional changes and immune challenge transcriptional changes than would be expected by chance.

To characterize the *Gba1b* mutant immune response in more detail, we investigated whether the transcriptional changes in *Gba1b* mutants resembled responses to any specific type of pathogen. In particular, as GlcCer is a known fungal virulence factor [32], we were interested in whether the immune response resembled the response to fungal infection. To do this, we compared our findings to data mined from previously published *Drosophila* RNA-Seq and microarray studies of immune response. Our data sources included sets of transcriptional changes detected in response to fungi, bacteria, viruses, and parasitoids [33–38]. We also compared our RNA-Seq data to a set of 62 core genes responsive to immune challenge, derived from a meta-analysis of 12 pathogen challenge studies [28]. See Methods and Fig S1 for details of our analyses. We found that transcripts from all the immune challenge lists were overrepresented among the transcripts increased in abundance in *Gba1b* mutants (∼7-fold to 18-fold enrichment, *p* < 0.005 by Fisher exact test; Fig 1D). The strongest enrichment was for gene expression associated with parasitoid attack. We then compared the RNA-Seq findings to the results from our previous proteomic analyses of *Gba1b* mutant heads [39]. To do this, we converted the previously mentioned lists of transcripts from immune response studies to lists of the proteins they encoded, and compared the converted lists to our head proteomics data. As in the RNA-Seq, proteins from all of the immune challenge lists were more likely to be increased in abundance in *Gba1b* mutants than would be expected by chance (Fig 1E). *Gba1b* mutants thus demonstrated a generalized immune response at the translational as well as the transcriptional level.

### Genetic manipulation of humoral immune factors does not rescue *Gba1b* mutant pathology

Given the many humoral factors increased in abundance in our RNA-Seq and proteomic data, we hypothesized that suppression of humoral immune pathways would ameliorate *Gba1b* mutant phenotypes. As in our previous work, our primary outcome measure was accumulation of insoluble ubiquitinated proteins in the head, which is a hallmark of neurodegenerative diseases [40–42]. For each manipulation, we measured insoluble ubiquitinated protein in the following four groups of flies:

*Gba1b* mutants with GAL4 driver and RNAi transgene

(tests the influence of decreasing expression of the gene of interest on *Gba1b* mutants)

*Gba1b* mutants with GAL4 driver and control RNAi transgene (e.g., *mCherry* RNAi)

(shows the phenotype of *Gba1b* mutants without intervention)

*Gba1b* controls with GAL4 driver and RNAi transgene

(shows any effect of the genetic manipulation in a wild-type background)

*Gba1b* controls with GAL4 driver and control RNAi transgene

(shows the phenotype of healthy flies; controls for possible effects of genetic background)

In most cases, the genetic manipulations showed no effect on control flies, we therefore report the results for *Gba1b* mutants with and without the intervention.

Recent reports have emphasized the importance of *Gba1b* in glia [43, 44], which are the primary immune-reactive cells of the nervous system [45], and we therefore first tested knockdown of humoral immune function in glia. We used the UAS/GAL4 tissue-specific expression system (pan-glial driver *repo-GAL4*) to knock down expression of transcription factors from the major humoral innate immune pathways [46–49]: *Rel* (Imd), *Dif* and *dl* (Toll), *Stat92E* (JAK/STAT), *kay* (JNK), and *Atf-2* (p38/MAPK). None of these manipulations had a significant effect on ubiquitinated protein accumulation in *Gba1b* mutants (Fig 2A). We therefore examined the effects of immune factor knockdown in fat body, the primary peripheral source of humoral immune effectors [50]. Knockdown of *Relish* using the adult fat body driver *ppl-GAL4* reduced ubiquitinated protein aggregates in the heads of both *Gba1b* mutants and controls (Fig 2B, Fig S2A-B), consistent with the previously reported influence of *Relish* on neurodegeneration [51, 52]. Further perturbations of the Imd pathway (a null mutant for *PGRP-LC* and the hypomorph *imd^1^*), however, showed no genetic interaction with *Gba1b* and did not alter insoluble Ub accumulation (Fig S2C-D). Fat body knockdown of the other immune-related transcription factors had no significant effect on Ub accumulation (Fig 2B).

**Figure 2:**
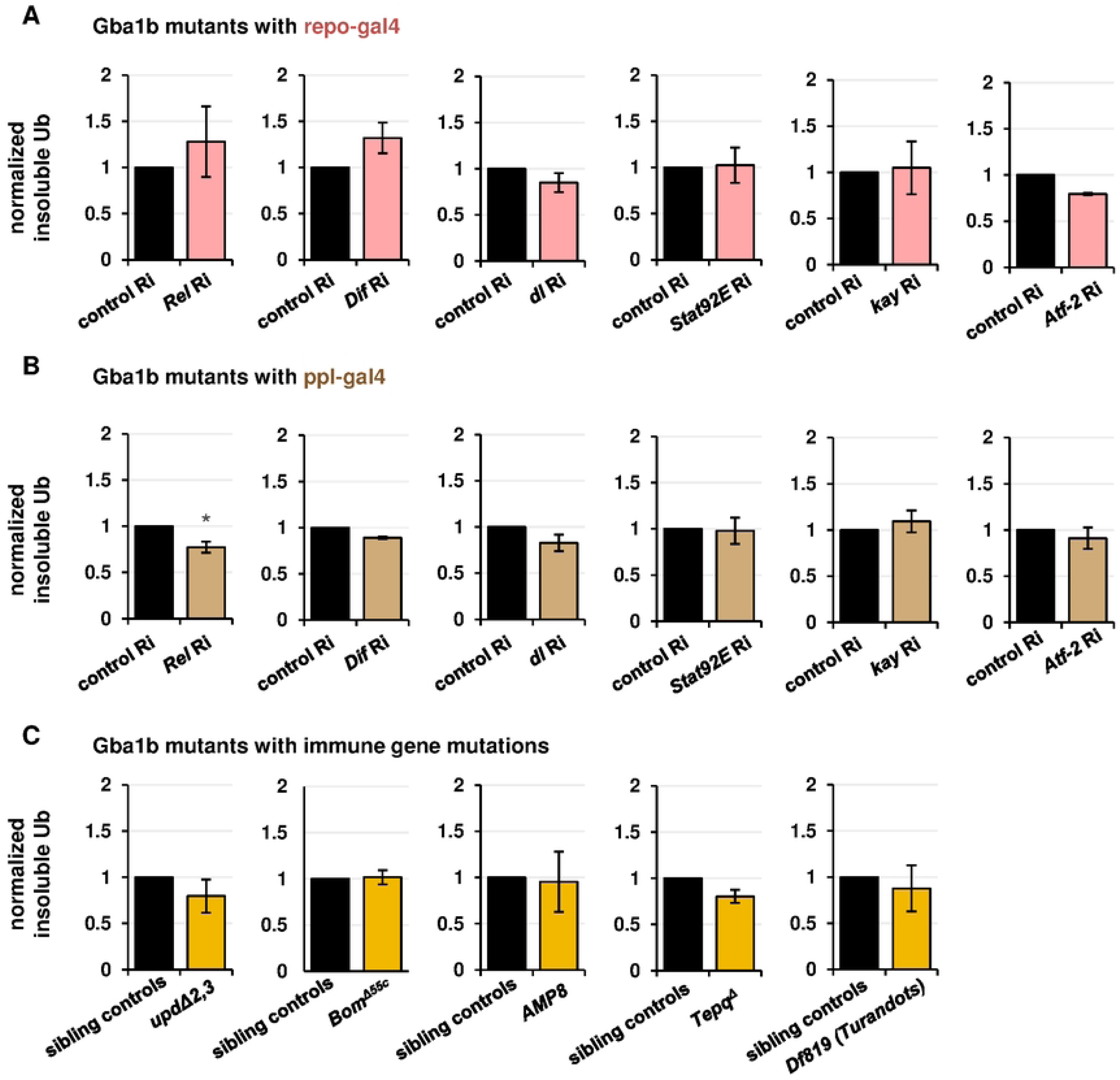
Suppression of humoral immune factors does not reduce accumulation of insoluble ubiquitinated proteins in *Gba1b* mutants. All panels represent quantification of anti-Ub (anti-ubiquitin) immunoblots using Triton-insoluble head samples from 10-day-old flies. Each graph compares *Gba1b* mutants with a genetic intervention to *Gba1b* mutants without the intervention. (A) RNAi to the transcription factor genes indicated, driven in glia with *repo-GAL4*. (B) RNAi to the transcription factor genes indicated, driven in fat body with *ppl-GAL4*. *Gba1b* mutants bearing mutations in immune genes. The first and last panels in this section represent immunoblots of whole bodies and the remainder represent immunoblots of heads.

As interfering with humoral immune expression in glia or fat body did not ameliorate Ub accumulation, we attempted to influence this phenotype using whole-organism gene ablation of immune effector groups. The alleles used were as follows (Fig 2C): a deletion of *Upd2* and *Upd3*, a deficiency removing 10 of 12 *Bomanin* genes, multiple deletions removing all known antimicrobial peptide genes on the second chromosome (*Def*, *AttA*, *AttB*, *AttC*, *Dro*, *Mtk*, *DptA*, and *DptB*), and a deletion of four thioester-containing proteins (*Tepq^Δ^*). None of these four interventions rescued the accumulation of ubiquitinated protein in *Gba1b* mutants. Finally, because we found prominent upregulation of *Turandot* family proteins in the RNA-Seq and proteomics, we tested for involvement of that family in the *Gba1b* mutant phenotype.

Heterozygous ablation of *TotA*, *TotB*, *TotC*, and *TotZ* using a large deficiency produced no effect on ubiquitin accumulation in the mutants (Fig 2C). We thus found no evidence to implicate canonical humoral immune pathways in the development of *Gba1b* mutant neuropathology.

### RNA-Seq and proteomics suggest prominent cellular immune activation in *Gba1b* mutants

Because the findings above indicated that humoral immune activation was not critically involved in *Gba1b* mutant neurological phenotypes, we considered the possibility that cellular rather than humoral immunity was involved. While *Drosophila* immune cells, or macrophages, were previously divided into three categories, recent single-cell RNA-Seq studies have revealed that they exist in a wide spectrum of states, from quiescent or proliferative states to states reflecting immune activation [53–58]. Using the classifications of Tattikota et al. and Cattenoz et al. [53, 54], we compiled a list of markers robustly associated with immune-activated macrophage subtypes in scRNA-seq studies. We also drew markers from a microarray study of mutants with excess lamellocytes, another activated immune cell type [59]. Although lamellocytes were previously considered to exist only in wasp-infested larvae [58, 60, 61], recent work has made clear that cells bearing lamellocyte biochemical markers can be found in adults [58], and we found that these markers were increased in abundance in *Gba1b* mutants. Markers of activated macrophages and lamellocytes were strikingly overrepresented among the transcripts increased in abundance in *Gba1b* mutants (Fig 3A), and the corresponding proteins were overrepresented as well (Fig 3B). We found the same increased enrichment when we compared our data to a proteomic analysis of lamellocytes (Fig 3B) [62]. The data as a whole strongly suggested increased activation of immune cells in *Gba1b* mutants.

**Figure 3.**
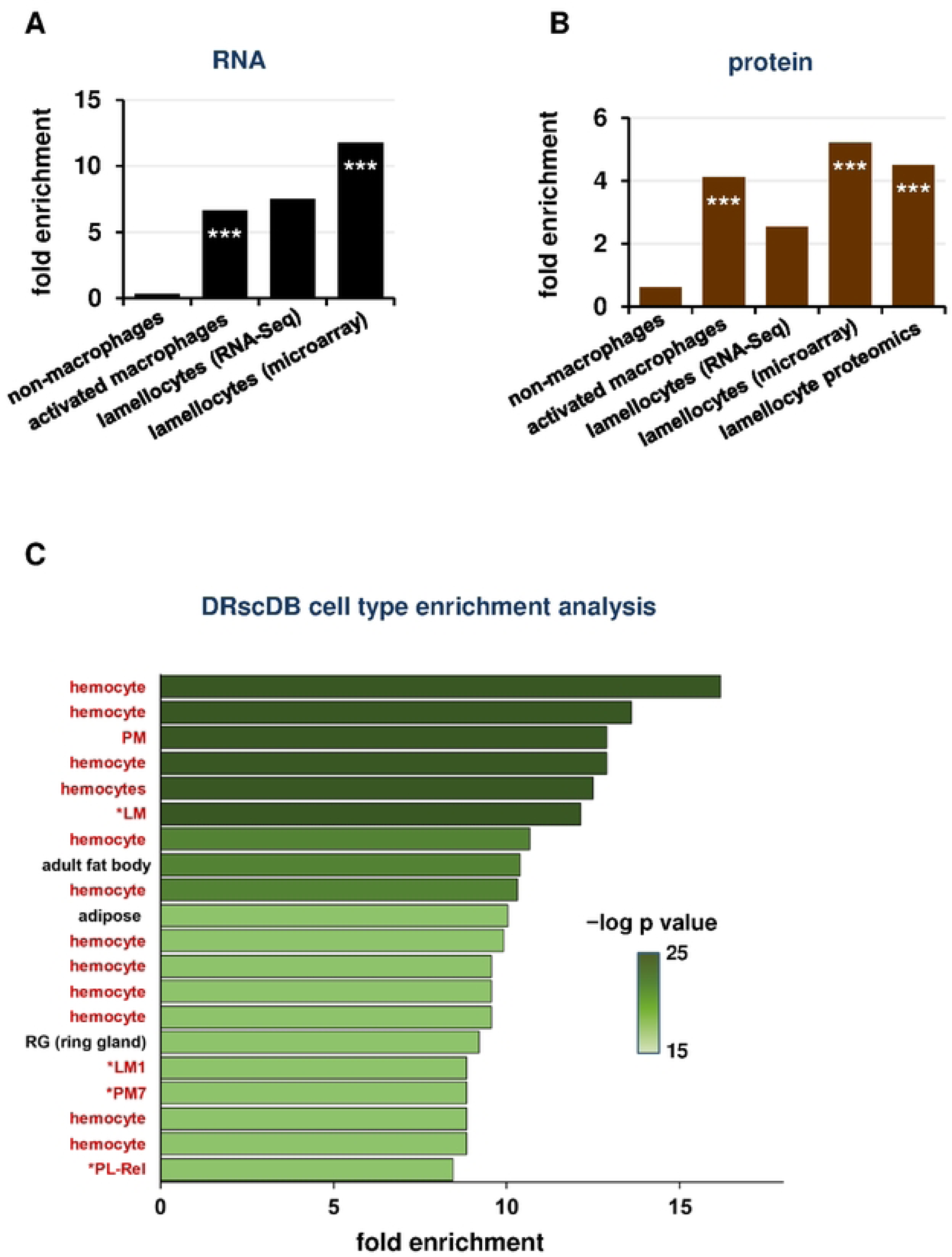
Analysis of RNA-Seq and proteomic data reveals increased abundance of markers associated with activated macrophage subtypes. (A-B) Enrichment of macrophage cell type markers in *Gba1b* mutant RNA-Seq (A) and proteomic data (B), calculated as in Figure 1. \*\*\**p* < 0.005 by Fisher exact test, indicating a significantly greater correspondence between *Gba1b* mutant transcriptional changes and activated immune cell marker lists than would be expected by chance. (C) Analysis of *Gba1b* RNA-Seq data using the Drosophila RNAi Screening Center (DRscDB) enrichment function. The top 20 terms by fold enrichment are shown. Terms indicating macrophages or macrophage subtypes (17 out of 20 cell types) are highlighted in red.

We further tested this theory using a single-cell RNA-Seq data resource from the Drosophila RNAi Screening Center [63]. The Enrichment function allows the user to enter a list of genes and compares the list to cell type markers from many single-cell RNA-Seq datasets. The user receives information on which cell types most closely match the set of genes submitted.

When we entered the genes significantly increased in abundance in *Gba1b* mutants, the cell types with which they were associated were overwhelmingly macrophages (here described with the invertebrate-specific term “hemocytes”), particularly activated macrophage subtypes (LM, PM7, PL-Rel; Fig 3C) [53–55]. This alternative analysis supported the idea that *Gba1b* mutant macrophages were abnormally activated.

### Fluorescent cell type markers show macrophage activation in *Gba1b* mutants

Having found evidence of macrophage activation in *Gba1b* mutants, we tested whether the activation could be detected on direct visualization of affected cells. We first tested whether there was a general proliferation of blood cells using the engineered fluorescent marker *SrpHemo-mCherry* [64], which is recognized as common to 99% of fly immune cells and maintains expression well into adulthood (Fig S3) [58]. Immunoblotting revealed no significant difference in marker abundance between *Gba1b* mutants and controls, and microscopy also revealed no obvious difference in total cell number (Fig 4A-C).

**Figure 4.**
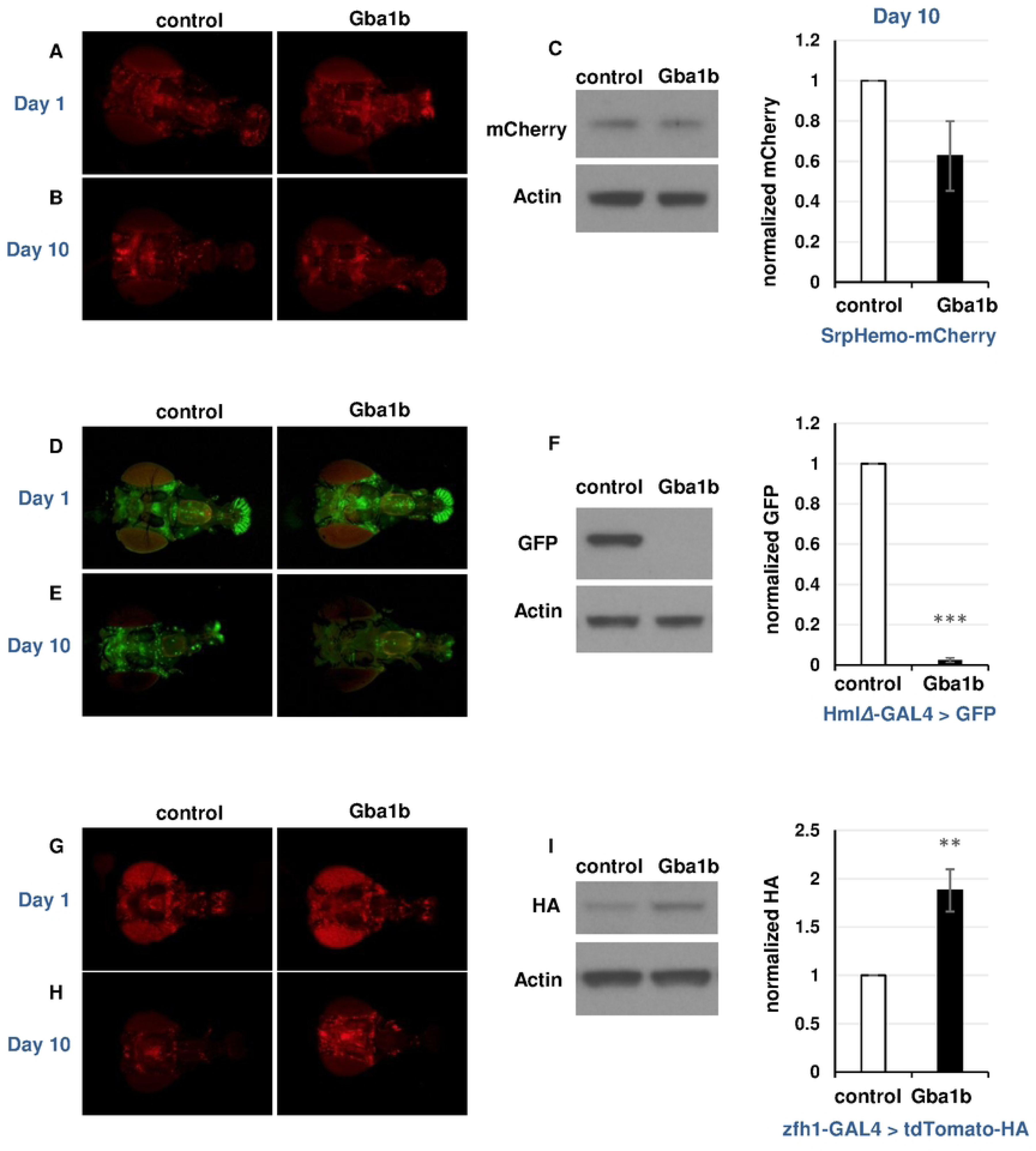
*In vivo* fluorescent cell type markers confirm an increased percentage of activated macrophages in *Gba1b* mutants. (A) *srpHemo-mCherry* fluorescence in heads from *Gba1b* mutants and control flies on day 1 of adult life. (B) *srpHemo-mCherry* fluorescence in heads at day 10. (C) Immunoblot and quantification of mCherry on day 10 heads from *Gba1b* mutants and controls expressing *srpHemo-mCherry*. (D) *HmlΔ-GAL4 > GFP* fluorescence in heads from *Gba1b* mutants and control flies on day 1 of adult life. (E) *HmlΔ-GAL4 > GFP* fluorescence in heads at day 10. (F) Immunoblot and quantification of GFP on day 10 heads from *Gba1b* mutants and controls expressing *HmlΔ-GAL4 > GFP*. (G) *zfh1-GAL4 > tdTomato-HA* fluorescence in heads from *Gba1b* mutants and control flies on day 1 of adult life. H) *zfh1-GAL4 > tdTomato-HA* fluorescence in heads at day 10. I) Anti-HA immunoblot and quantification on day 10 heads from *Gba1b* mutants and controls expressing *zfh1-GAL4 > tdTomato-HA*. All protein samples were extracted using RIPA buffer as described in Materials and Methods. Heads were imaged caudal side down, with the ventral aspect to the left. Significance was tested using Student’s *t* test, ** = *p* < 0.01, *** = *p* < 0.005.

Given that there was no gross change in macrophage number, we hypothesized that *Gba1b* mutants had a shift in the relative abundance of macrophage subpopulations. We decided to examine macrophage subpopulations in *Gba1b* mutants using several GAL4 drivers and markers. Because a number of macrophage GAL4 drivers stop driving expression early in adult life [65], we screened available reagents for GAL4 drivers and markers that continued to drive expression in macrophages 10 days after the fly reached adulthood. We found two relevant GAL4 drivers: *HmlΔ-GAL4* and *GMR35H09-GAL4* (Fig S3). *GMR35H09-GAL4* contains DNA sequences derived from the *zfh1* gene [66], which is highly expressed in activated macrophage subtypes [53, 54], and we therefore refer to it henceforth as *zfh1-GAL4*.

Using these tools, we tested whether the evidence of immune cell activation seen on RNA-Seq could be detected via expression of fluorescent markers. We predicted that markers of activated cell types would be increased in *Gba1b* mutants relative to controls, while other macrophage types would be decreased. We tested one marker known to decrease in activated macrophages (*HmlΔ-GAL4 > UAS-GFP*, henceforth *HmlΔ-GFP*) [64, 67] and one associated with activated macrophages (*zfh1-GAL4* driving *tdTomato-HA*) [53, 54]. In one-day-old flies, mutants and controls showed no difference in fluorescent signal from these markers, consistent with the lack of obvious phenotypes in *Gba1b* mutants at that age (Fig 4D, 4G) [19]. However, at 10 days of age, immunoblotting and microscopy showed that *HmlΔ-GFP* signal was severely depleted (Fig 4E-F); the same result was found with *HmlΔ-dsRed* (Fig S4A), while *tdTom-HA* driven by *zfh1-G4* was increased in abundance in *Gba1b* mutants vs. controls (Fig 4H-I). These findings indicated that the macrophage population in *Gba1b* mutants is shifted toward activated cell types. We further tested this hypothesis using *VT17559-GAL4,* whose promoter comes from the activated macrophage marker *Lis-1* [65]. When we drove *tdTom-HA* with this driver, as with *zfh1-GAL4*, signal was increased in *Gba1b* mutant macrophages at day 10 (Fig S4B). Together, these findings demonstrate age-dependent cellular immune activation in *Gba1b* mutants.

### Restoring GCase activity to activated macrophages rescues *Gba1b* mutant phenotypes

To identify the contribution of immune cell activation to pathogenesis in *Gba1b* mutants, we tested whether driving expression of *Gba1b* in activated macrophages would ameliorate the mutants’ phenotypes, including macrophage activation, insoluble Ub accumulation, short lifespan, and impaired locomotor performance. We hypothesized that driving *Gba1b* in the macrophages showing aberrant activation in the mutants would rescue their phenotypes, while driving *Gba1b* in other macrophage subtypes would not. To test this, we used three of the GAL4 drivers described above: *Srp-GAL4*, which drives in most macrophage subtypes; *zfh1-GAL4*, which drives in the aberrantly activated macrophages; and *eater-GAL4*, which drives in a different subset of macrophages [54]. We predicted that *Srp-GAL4* and *zfh1-GAL4* would rescue *Gba1b* mutant phenotypes and that *eater-GAL4* would not. When we examined the accumulation of insoluble ubiquitinated protein in *Gba1b* mutants, our predictions were validated (Fig 5A-C).

**Figure 5.**
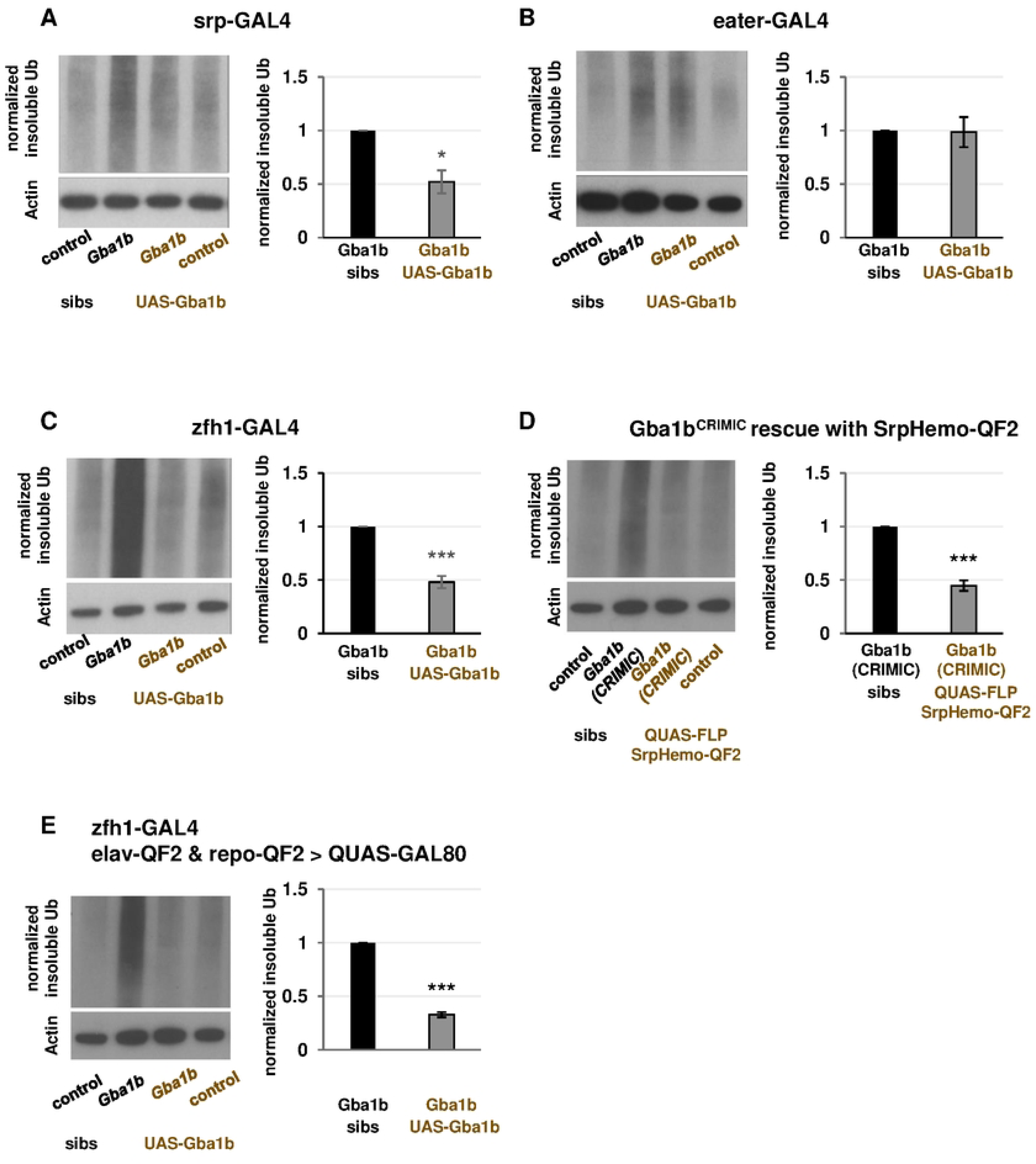
Expression of *Gba1b* in activated macrophages rescues insoluble ubiquitin accumulation in *Gba1b* mutants. (A-C) Immunoblot and quantification of insoluble ubiquitinated head proteins from *Gba1b* mutants and controls with the following genetic manipulations: (A) *UAS-Gba1b* under the control of the pan-macrophage driver *Srp-GAL4*. (B) *UAS-Gba1b* under the control of the driver *eater-GAL4*, which drives in a subset of macrophages. (C) *UAS-Gba1b* under the control of the driver *zfh1-GAL4*, which drives in a subset of activated macrophages. (D) Immunoblot and quantification of insoluble ubiquitinated head proteins from flies with endogenous *Gba1b* gene expression restored in macrophages. This was achieved by excising the transposon causing *Gba1b* loss of function from the *Gba1b*^CRIMIC^ allele (expressing FLP recombinase under the control of *SrpHemo-QF2*). (E) Immunoblot and quantification of insoluble ubiquitinated head proteins from flies expressing GAL80 under the control of pan-neuronal driver *elav-QF2* and pan-glial driver *repo-QF2*, as well as *Gba1b* under the control of the driver *zfh1-GAL4.* Significance was tested using Student’s *t* test, * = *p* < 0.05, *** = *p* < 0.005. “Sibs” refers to siblings of the flies that bear both a UAS construct and a GAL4 driver. Siblings may have the GAL4 or the UAS, but not both.

We tested alternate explanations for our findings. First, we addressed the possibility that rescue of insoluble ubiquitinated protein levels by *Gba1b* expression in macrophages was the result of supraphysiological gene expression produced by the UAS-GAL4 system. To do this, we used flies bearing the *Gba1b*^CRIMIC^ allele, which disrupts *Gba1b* function [43], and removed the transposon to restore function of the endogenous gene. Because no *zfh1-QF2* driver was available, we used the pan-macrophage driver *SrpHemo-QF2* to target transposon removal to macrophages. This manipulation robustly rescued the accumulation of insoluble ubiquitinated proteins in *Gba1b* mutant heads, indicating that our findings were not an artifact of high expression levels (Fig 5D). Second, to address the possibility that our findings were the result of *Gba1b* misexpression in neurons or glia, we created flies that expressed *Gba1b* in macrophages but had a blockade of neuronal and glial GAL4 activity. Specifically, we created a strain expressing the GAL4 repressor GAL80 under the control of both *elav-QF2* and *repo-QF2*, drivers that do not interact with the UAS-GAL4 system [68]. In these flies, we drove *Gba1b* using *zfh1-GAL4* as above. We found that rescue of ubiquitin accumulation in these animals was comparable to the rescue in flies without GAL80 inhibition, indicating that misexpression in neurons or glia does not explain our results (Fig 5E).

We next tested whether *Gba1b* expression under control of *zfh1-GAL4* also ameliorated other phenotypes of *Gba1b* mutants, including macrophage activation, reduced lifespan, locomotor deficits, and autonomic nervous system abnormalities. To test macrophage activation, we expressed *tdTomato* under the control of *zfh1-lexA* as a measure of fluorescence that would not be affected by GAL4 drivers [69, 70]. We examined this marker in *Gba1b* mutants and controls with and without *zfh1-GAL4*–driven restoration of GCase activity. Restoring GCase activity in *Gba1b* mutants caused a substantial decrease in fluorescent signal, indicating a normalization of the proportion of activated macrophages (Fig 6A). As macrophage activation in *Gba1b* mutants is also marked by loss of *HmlΔ-dsRed* (Fig S4A), we tested whether driving *Gba1b* with *zfh1-GAL4* would reverse this phenotype as well, and we found that this was the case (Fig S5). These findings show that macrophage activation in *Gba1b* mutants is dependent on loss of GCase activity in macrophages. We then tested whether *Gba1b* expression in activated macrophages would also rescue the mutants’ deficits in lifespan, locomotor performance, and enteric nervous system function. Expressing *Gba1b* in activated macrophages ameliorated the mutants’ shortened lifespan (Fig S6) and restored their climbing ability as measured by the RING assay [71](Fig 6B). We also assayed gut transit time, a measure of enteric nervous system function, using a procedure modified from Olsen and Feany (Fig S7A) [72]. *Gba1b* mutants, like some other neurodegeneration model flies, display markedly impaired clearance of food from the gut (Fig S7B-C). Restoring *Gba1b* to activated macrophages completely normalized gut transit (Fig 6C).

**Figure 6.**
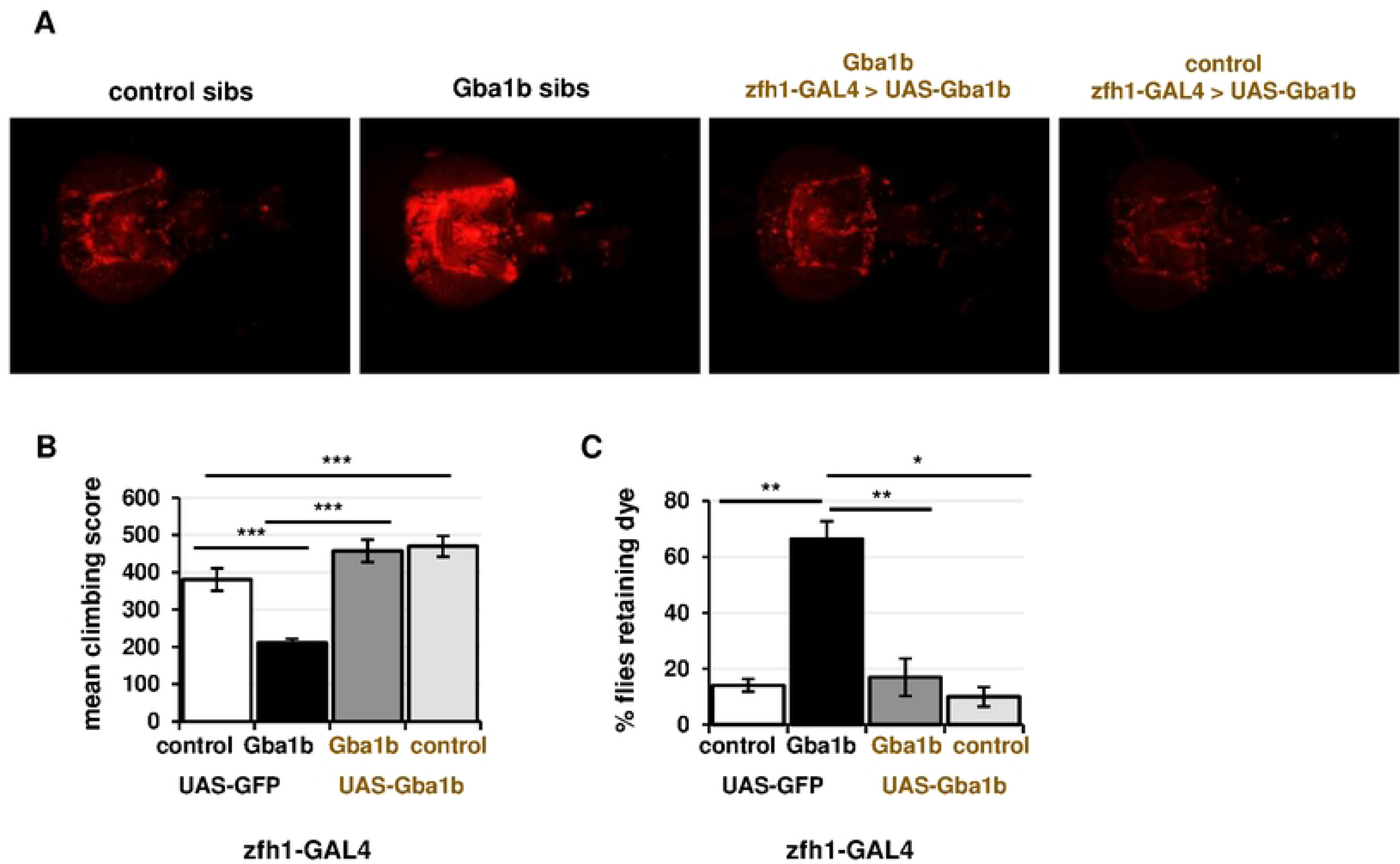
Restoring GCase function in activated macrophages rescues *Gba1b* mutant phenotypes. (A) Visualization of macrophage activation (using *zfh1-lexA > tdTom*) in 10-day-old *Gba1b* mutants and controls with or without rescue expression of *Gba1b* under the control of *zfh1-GAL4* (*n* = 6-8 animals per genotype). (B) RING climbing assay in *Gba1b* mutants and controls with or without rescue expression of *Gba1b* under control of *zfh1-GAL4*. All flies were 13-day-old females. \*\**p* < 0.01 (C) Gut transit assay, adapted from Olsen and Feany [72], to measure enteric nervous system function. Fifteen-day-old female flies (*Gba1b* mutants and controls with or without *zfh1-GAL4* driving *UAS-Gba1b)* were scored for the presence of blue dye in the abdomen 2.5 h after being transferred from dyed to plain food.

### Macrophage activation appears to be caused by cell-autonomous accumulation of GlcCer

Having shown that cellular immune activation is important to the development of *Gba1b* mutant phenotypes, we wanted to test the mechanism by which they are activated. *Gba1b* mutants have been shown to have compromised circadian rhythms due to lipid alterations in central clock neurons [44], and disruption of circadian rhythm can lead to abnormal macrophage activation [73, 74]. We therefore considered the possibility that normalizing the activity of central clock neurons in *Gba1b* mutants might be sufficient to ameliorate both Ub accumulation and macrophage activation. We expressed *Gba1b* with *Pdf-GAL4*, which has been shown to rescue cyclic neurite remodeling in sLNv clock neurons [44]. We assessed macrophage activation as described above (*zfh1-lexA* driving *tdTom* and *Pdf-GAL4* driving *UAS-Gba1b*). Restoring *Gba1b* function to *Pdf-GAL4* neurons did not ameliorate macrophage activation (Fig S8) or significantly reduce Ub accumulation (Fig 7A). These findings suggest that GlcCer accumulation in sLNv clock neurons is not a major cause of ubiquitinated protein accumulation in *Gba1b* mutant heads and is not responsible for macrophage activation.

**Figure 7:**
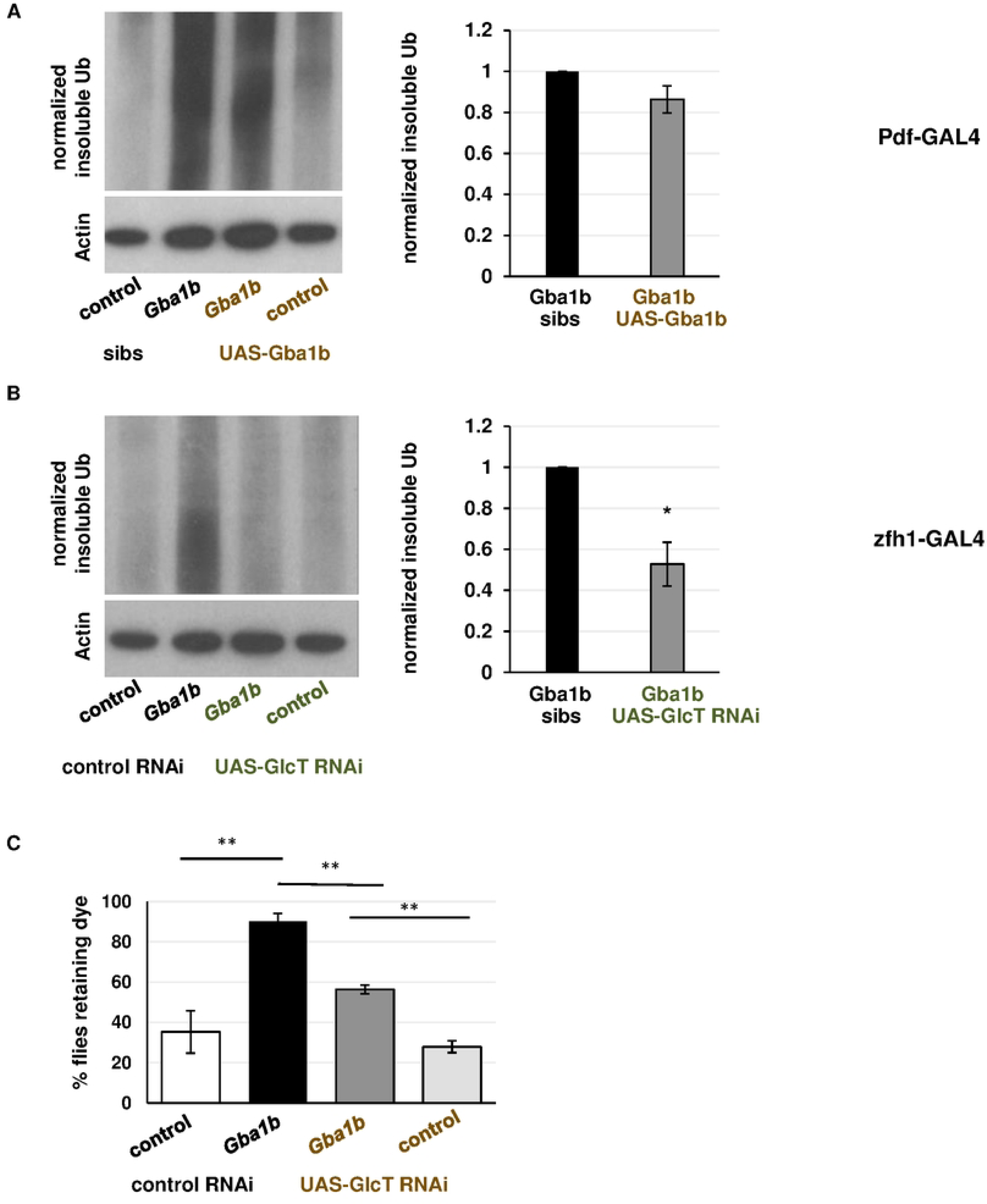
GlcCer in activated macrophages is critical to the development of protein aggregation in *Gba1b* mutants. (A-B) Immunoblot and quantification of insoluble ubiquitinated head proteins from flies expressing (A) *Gba1b* under the control of *Pdf-GAL4*, which drives in circadian rhythm neurons or (B) *GlcT RNAi* under the control of the activated hemocyte driver *zfh1-GAL4*. C) Gut transit assay on *Gba1b* mutants with or without *zfh1-GAL4* driving *UAS-GlcT*. Female flies, 20-22 days old, were tested as described above. \**p* < 0.05, \*\**p* < 0.01, \*\*\**p* < 0.005 by Student’s *t* test.

Given the above findings, we hypothesized that GlcCer accumulation in the macrophages themselves was responsible for the aberrant activation. We reduced the level of GlcCer in activated macrophages using knockdown of the synthetic enzyme for GlcCer, *GlcT* [75], under control of *zfh1-GAL4.* This manipulation reduced the accumulation of insoluble Ub in *Gba1b* mutant heads (Fig 7B). As noted above, restoring endogenous levels of *Gba1b* to macrophages also ameliorated this phenotype, suggesting that GlcCer levels in macrophages are critical for the development of ubiquitinated protein accumulation.

Finally, we tested the functional significance of GlcCer levels in the activated macrophages and found that GlcT knockdown improved autonomous nervous system function (Fig 7C). Effects on climbing could not be assessed because *zfh1-GAL4* knockdown of GlcT impaired locomotion even in control flies (Fig S9). We conclude that excess GlcCer in macrophages promotes aberrant activation in a cell-autonomous manner, and that these activated macrophages promote the development of behavioral and biochemical abnormalities associated with GCase deficiency.

## Discussion

In this work, we have shown that the development of central nervous system pathology in GCase-deficient flies requires activation of peripheral immune cells. This activation appears to be a direct, cell-autonomous consequence of membrane lipid alterations in macrophages due to excess GlcCer.

Humoral immunity was also clearly activated in *Gba1b* mutants, but a large set of genetic manipulations found no evidence that the humoral immune system was involved in the development of neurological phenotypes. The most likely explanation for this is that the increased production of humoral factors is a consequence of macrophage activation [76–78].

Some previously published models of neurodegenerative disease have shown that excess production of humoral immune factors is sufficient to cause neurodegeneration [51, 52, 79, 80] and in *Drosophila*, the detrimental effects of humoral activation were blocked by interfering with the upstream immune transcription factor *Relish* [51, 52]. Other work, by contrast, has shown that increased abundance of antimicrobial peptides promoted survival and recovery after head injury [81]. In our model, modifying humoral immune function appeared neither helpful nor harmful. Our results underscore the fact that, while neuroinflammation is a common finding in neurodegenerative states [82], it is necessary to test the functional significance of immune changes before concluding that they are pathogenic [81, 83].

Inappropriate macrophage activation and even central nervous system invasion has been described in various neurodegenerative diseases [84–87], but reports conflict as to whether this phenomenon promotes or opposes neurodegeneration [88–93]. Similarly, while macrophage alterations are a hallmark finding in severe GCase deficiency [13, 14], their role in neurological abnormalities has not been established. Earlier reports emphasized the importance of lipid abnormalities in neurons [94], although these have not been universally found. Two recent papers indicated that *Gba1b* acts primarily in glia, which receive and degrade the GlcCer produced in neurons [43, 44]. In vertebrates, loss of GCase function in microglia, an immune-specific glial subtype, has been implicated in neuronal damage [95]. In *Drosophila*, the functions associated with vertebrate microglia are distributed among multiple types of glia [96]. While restoring GCase function in glia did rescue *Gba1b* mutant phenotypes [43, 44], we found that expressing *Gba1b* in activated macrophages rescued ubiquitin-protein accumulation even when GAL4-driven *Gba1b* expression was blocked in both neurons and glia. Our results thus point to the importance of peripheral as well as central immune cells in the development of neuropathology. This finding raises two major questions: how does GlcCer accumulation lead to aberrant macrophage activation, and how does aberrant macrophage activation cause *Gba1b* mutant phenotypes?

GCase deficiency could lead to macrophage activation in several ways. The mechanism best supported by our data is that GlcCer accumulation within macrophages causes cell-autonomous activation by mimicking the changes in lipid balance that normally occur during macrophage activation. First, the process of macrophage activation involves synthesis of sphingolipids and a large increase in cellular sphingolipid content [97], and other factors that change lipid balance (e.g., high-fat diet) have been shown to cause pro-inflammatory macrophage phenotypes [98, 99]. Second, reducing GlcCer synthesis within activated macrophages was sufficient both to reduce ubiquitin-protein aggregation and to ameliorate gut transit abnormalities. Third, humoral factors are normally involved in the signaling that leads to macrophage activation, but reduction of humoral factors had no effect on *Gba1b* mutant phenotypes [100]. Alternatively, GlcCer accumulation in macrophages could cause hyperreactivity to normal immune stimuli, as previously reported in macrophages from PD patients with *GBA* mutations [15]. Atilano et al. [26] partially addressed this possibility by raising GCase-deficient flies in germ-free conditions. The improvements in lifespan and climbing after this intervention, however, were modest compared to those seen when we restored GCase activity in activated macrophages. Together, these findings suggest that macrophage activation is more a primary result of lipid changes than a sensitization to external immune challenges.

An equally important question is how macrophage activation contributes to the development of central nervous system abnormalities (Fig 8). First, macrophages could enter the brain and act directly on central nervous system cells. Invasion of the CNS by peripheral immune cells has been reported in multiple mammalian models of neurodegenerative disease [88, 101–103], and more recently in immune-challenged *Drosophila* [87]. In the *Drosophila* study, the invading macrophages caused direct destruction of neuronal processes through phagocytosis [87]. Invading macrophages can also cause harm through mechanisms such as signaling to microglia [104], oxidative stress [105], or glutamate excitotoxicity [106, 107]. However, not all models of neuropathology in GCase deficiency show blood-derived macrophages in the brain [108–110], and macrophages also influence brain function from beyond the blood-brain barrier [104]. Given our previous findings of altered extracellular vesicle biology in *Gba1b* mutants [39], long-distance signaling from macrophages to brain is a possible mechanism [104]. In a *Drosophila* Alzheimer’s disease model, macrophages clustering outside the brain promoted neurodegeneration via TNF-JNK signaling [92]. In addition, phagocytosis by macrophages can impact the central nervous system through its effect on the abundance of aggregation-prone proteins. Significant quantities of these proteins (including alpha-synuclein, tau, and amyloid beta) [111] are constantly released from the brain; in the case of amyloid beta, about half of the protein released from the brain is normally eliminated in the periphery [112]. Moreover, interfering with macrophage clearance of amyloid beta is reported to lead to increased levels of the protein in the brain [113]. One possible mechanism of harm, therefore, is inadequate peripheral clearance of aggregation-prone proteins by macrophages in *Gba1b* mutants, leading to accumulation of ubiquitinated protein in brain and periphery alike. As noted above, abnormalities in phagocytosis have been described in macrophages from GCase-deficient humans [18, 95]. Further investigation will be required to determine the relative contributions of these or other mechanisms of pathogenesis. Our findings do suggest, however, that targeting both peripheral and central immune cells may be necessary to prevent neurodegeneration caused by GCase deficiency.

**Figure 8:**
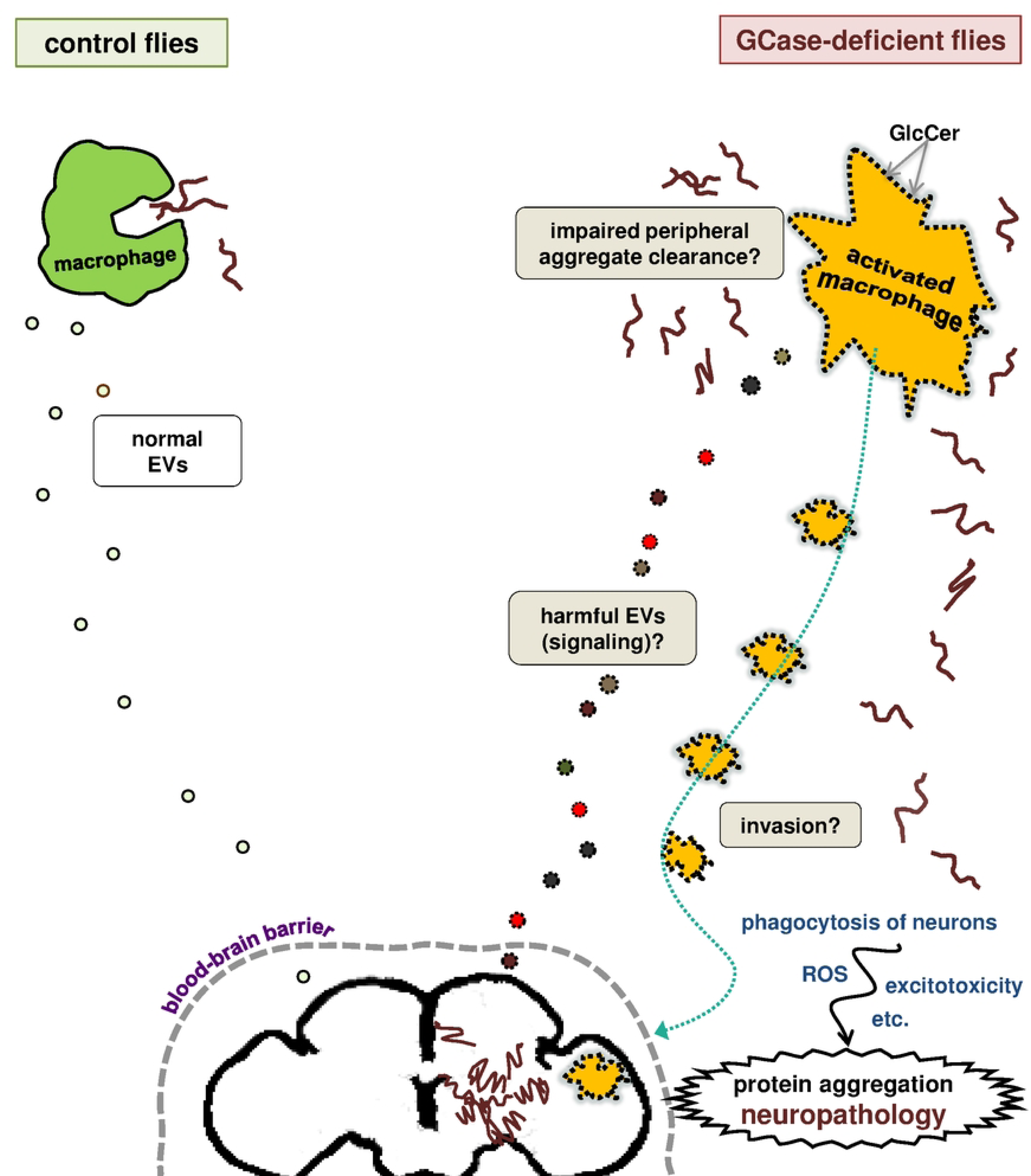
**Possible mechanisms of pathogenesis due to aberrant macrophage activation in *Gba1b* mutants**. Model: Excess GlcCer in macrophages causes activation via mimicry of the lipid changes associated with normal activation, with several possible detrimental outcomes that are not mutually exclusive. 1) Invasion: Activated macrophages may migrate into the brain or just outside the blood-brain barrier. They may cause damage by short-range mechanisms such as phagocytosis of neurons, release of ROS, or excitotoxicity. 2) Signaling: Activated macrophages may release harmful extracellular vesicles (EVs) or other signals that cause long-distance damage to the brain. 3) Failed peripheral aggregate clearance: Activated macrophages may have decreased capacity for phagocytosis, leading to inadequate clearance of aggregation-prone proteins from the periphery and their consequent accumulation in the brain.

## Acknowledgments

We thank Utpal Banerjee, Bruno Lemaitre, Iwan Evans, and Steven Wasserman for fly stocks and Chris Frazar of the University of Washington Northwest Genomics Center for sequencing services. Amy Platenkamp contributed to immune knockdown experiments.

## Materials and methods

### *Drosophila* strains and culture

Fly stocks were maintained on standard cornmeal-molasses food at 25°C on a 12:12 light/dark cycle. The *Gba1b* (*Gba1b^ΔTT^*), *Gba1b^rv^*, and *Gba1b^MB03039^* (*Gba1b Minos*) alleles have been previously described [19, 39]. “*Gba1b* mutants” refers to any combination of *Gba1b^ΔTT^* homozygotes, *Gba1b^ΔTT^/Gba1b^MB03039^* transheterozygotes, and *Gba1b^MB03039^* homozygotes.

These alleles have identical effects on the ubiquitinated protein accumulation phenotype [39]. Unless otherwise specified, controls for the genetic manipulation are siblings generated from the same cross and bear a balancer chromosome. In some cases, flies bearing a control transgene were used (*UAS-mCherry RNAi* for third chromosome transgenes, *UAS-lexA RNAi* for second chromosome transgenes). All third chromosome transgenes or alleles tested for modifying effects on *Gba1b* mutant phenotypes were recombined with the *Gba1b Minos* allele.

Endogenous *Gba1b* expression was restored using the following genotypes: 

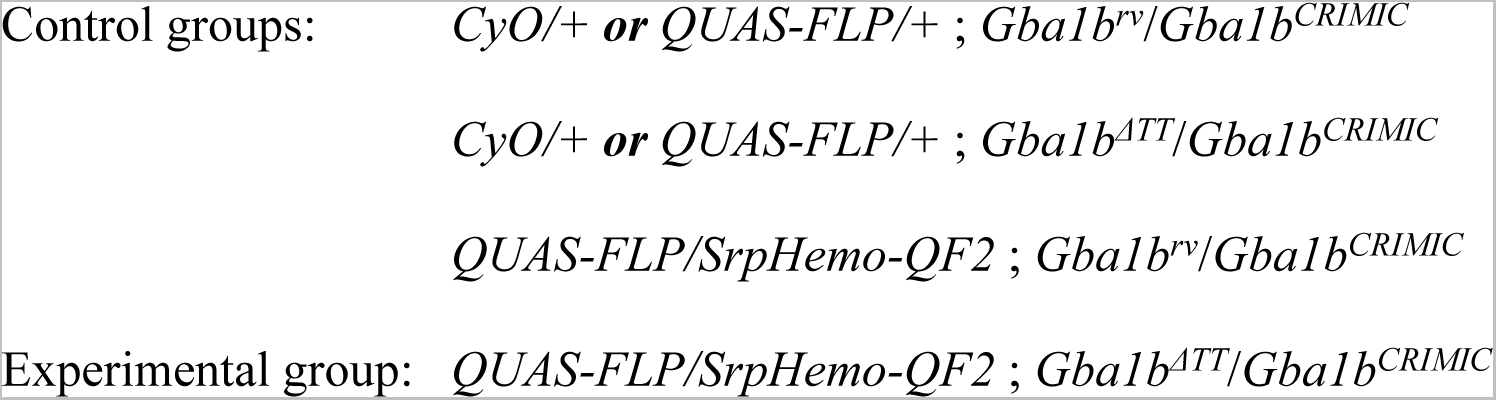

Other strains and alleles are as described in Table 1.

**Table 1.**
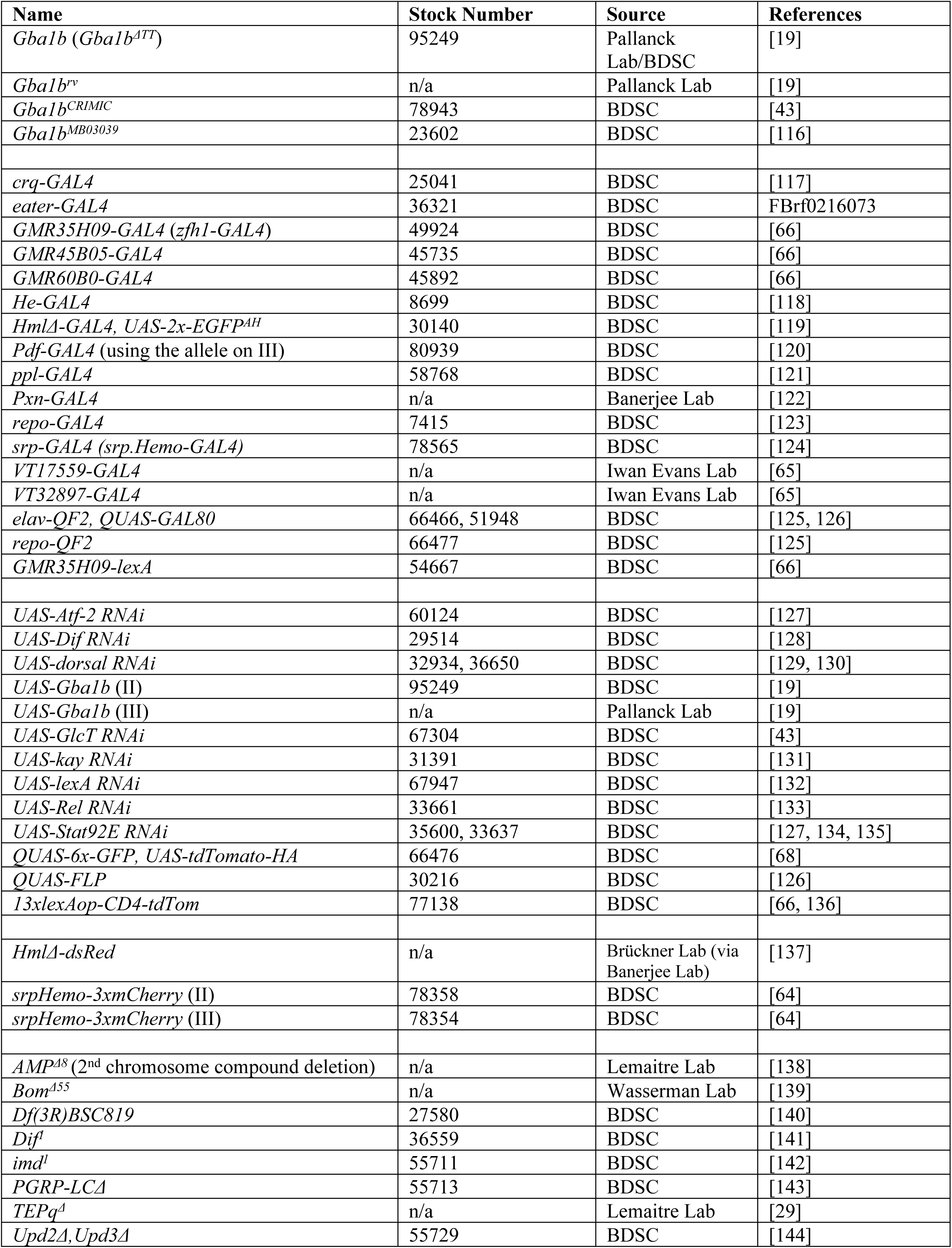
Fly stock information.

### RNA-Seq Analyses

#### RNA extraction

Heads from 11-day-old male *Gba1b* mutant and control flies were cut off using razor blades, on ice, and homogenized in TRIzol (500 μL per 50 heads). The homogenate was flash-frozen and stored at −80°C. RNA was extracted using a Direct-zol kit (Zymo Research R2050) according to the manufacturer’s instructions, using DNase treatment on column, and was stored at −80°C.

#### RNA sequencing

RNA sequencing was performed at the University of Washington Northwest Genomics Center (NWGC). RNA concentration was measured using the Quant-iT RNA assay (Invitrogen), and RNA integrity was tested using a fragment analyzer (Advanced Analytical). Total RNA was Total RNA is normalized to 12.5 ng/μL in a total volume of 47 μL on the Perkin Elmer Janus Workstation (Perkin Elmer, Janus II). Poly-A selection and cDNA synthesis were performed using the TruSeq Stranded mRNA kit as outlined by the manufacturer (Illumina RS-122-2103). All steps were automated on the Perkin Elmer Sciclone NGSx Workstation. Final RNA-Seq libraries were quantified using the Quant-it dsDNA High Sensitivity assay, and library insert size was checked using a fragment analyzer (Advanced Analytical; kit ID DNF474).

Samples in which adapter dimers constituted more than 3% of the electropherogram area were failed prior to sequencing. Technical controls (K562,Thermo Fisher Scientific AM7832) were compared to expected results to ensure that batch-to-batch variability was minimized. Successful libraries were normalized to 10 nM for submission to sequencing. Barcoded libraries were pooled using liquid handling robotics prior to loading. Massively parallel sequencing-by-synthesis with fluorescently labeled, reversibly terminating nucleotides was carried out on the MiSeq sequencer. Base calls were generated in real time on the MiSeq instrument, and demultiplexed, unaligned fastq files were generated by Samtools bcl2fastq.

#### GO Term Enrichment Analysis

GO Biological Process and KEGG pathway analyses were performed on RNA-Seq data using the PANGEA Gene Ontology tool [31] with Benjamini & Yekutieli correction for multiple testing. The cutoff for inclusion of terms or pathways was an adjusted *p* value < 0.05.

#### Cell type marker analysis using DRscDB

We used DRscDB, a manually curated single-cell RNA-Seq database [63], to identify the *Drosophila* cell types with markers most closely resembling the transcriptional changes in *Gba1b* mutants vs. controls. We performed this analysis using the Single Gene List Enrichment feature. All 379 transcripts increased in abundance were included. From the 102 cell marker terms with significant enrichment (Benjamini & Yekutieli correction), we selected the 20 cell types with the greatest fold enrichment.

### Proteomics

Proteomic studies of fly heads were performed as previously described [39].

### Comparisons of transcriptomic and proteomic data to published data

Our aim in these analyses was to characterize our *Gba1b* mutant findings by evaluating their resemblance to other sets of findings associated with immune activation. These included acute responses to immune stimuli and markers representative of activated immune cell types.

#### Creating marker lists

For each comparison dataset, we compiled a “marker list” of all transcripts with significantly increased abundance. For the immune challenge studies, this represented all transcripts increased in abundance after the relevant stimulus; for the immune cell type studies, the list was all transcripts enriched in the target cell type compared to other cell types studied. In some cases, the marker list was generated from two closely related datasets (e.g., transcripts increased in abundance in both of two parasitoid infestation studies). In one case, we used a proteomic dataset by converting the proteins into gene/transcript equivalents [62]. Each marker list was finalized by eliminating any transcripts not also detected in the *Gba1b* RNA-Seq dataset. Details of the studies, criteria used, and numbers of transcripts included are given in Table 2 for immune challenge studies and in Table 3 for cell type marker studies.

**Table 2.** Details of immune challenge comparison datasets.

| Class | Pathogen | Source | Criteria | Combination | # transcripts in list | # list transcripts detected in *Gba1b* | # ↑ in *Gba1b* | # proteins detected in *Gba1b* | # proteins ↑ in *Gba1b* |
| --- | --- | --- | --- | --- | --- | --- | --- | --- | --- |
| **Bacteria** | 10 species of bacteria | Troha et al. [33]: “core” bacterial response | ↑ in response to $\geq 7$ species | n/a | 166 | 159 | 54 | 68 | 30 |
| **Viruses** | Flock House, Sindbis, & C virus | Kemp et al. [34] | ↑ in response to all 3 viruses | n/a | 42 | 36 | 16 | 19 | 6 |
| **Fungi** | *B. bassiana* | Moskalev et al. [35], 100 cfu, time not given | Transcripts ↑ in abundance |  | 158 | 132 | 40 | 40 | 18 |
| | *E. muscae* | Elya et al. [36] | FC $\geq 1.5$ in whole flies 48 h after infection | Union (in either set) | | | | | |
| **Parasitoids** | *Ganaspis* sp. | Higareda Alvear et al. [37] | ↑ in abundance at 24 h |  | 40 | 31 | 21 | 12 | 5 |
|  | *A. tabida* | Salazar-Jaramillo et al. [38] | ↑ abundance, 5 h or 50 h | Intersection (in both sets) |  |  |  |  |  |
| **Immune meta-analysis** | “genes significantly induced by infection” | Waring et al. [28] | (selected by authors) | n/a | 62 | 55 | 28 | 23 | 10 |

**Table 3.** Details of activated immune cell comparison datasets.

| Cell Class | Source | Cutoffs | Combination | # transcripts, original list | # list transcripts detected in *Gba1b* | # increased in *Gba1b* | # proteins detected in *Gba1b* | # proteins increased in *Gba1b* |
| --- | --- | --- | --- | --- | --- | --- | --- | --- |
| **Non-hemocytes** | Tattikota et al. [53] | all | n/a | 656 | 642 | 8 | 333 | 18 |
| **Activated macrophages** | Tattikota et al. [53]: “activated hemocytes” as defined by authors (PM3, PM4, PM5, PM6, PM7) | $\log_2\text{FC} \geq 0.5$ vs. other subtypes, adjusted $p \leq 0.05$ | | | | | | |
| | Cattenoz et al. [54] PL-AMP, PL-vir1, PL-Rel | $\text{FC} \geq 2$ vs. other subtypes, adjusted $p \leq 0.05$ | Union (in either set) | 195 | 190 | 44 | 85 | 30 |
| **Lamellocytes (RNA-Seq)** | Tattikota et al. [53]: LM1, LM2 | $\log_2\text{FC} \geq 0.5$ vs. other subtypes, adjusted $p \leq 0.05$ | | | | | | |
|  | Cattenoz et al. [54]: LM1, LM2 | all | Intersection (in both sets) | 46 | 42 | 11 | 32 | 7 |
| **Lamellocytes (microarray)** | Kwon et al. [59] *hop<sup>tum</sup>* mutants | $\text{FC} \geq 2.5$ and adjusted $p \leq 0.05$ | | | | | | |
| | Kwon et al. [59] *Nurf301* mutants | $\text{FC} \geq 2.5$ and adjusted $p \leq 0.05$ | Intersection (in both sets) | 119 | 108 | 46 | 44 | 17 |
| **Lamellocytes (proteomics)** | Wan et al. [62] | 2D proteomics, emPAI > 1 | n/a | n/a | n/a | n/a | 62 | 24 |

#### *Comparing marker lists to* Gba1b *RNA-Seq data*

For each marker list, we determined the number of transcripts that were also increased in abundance in *Gba1b* mutants. We divided that value by the *whole-dataset* percentage of transcripts with increased abundance in *Gba1b* mutants, which was 3.48% for RNA-Seq and 8.58% for proteomics. The result was an enrichment score describing the degree of overlap between the marker list and the set of transcripts increased in abundance in *Gba1b* mutants. We determined whether that overlap was greater than would be predicted by chance alone using Fisher’s exact test.

#### *Comparing marker lists to* Gba1b *proteomic data*

We compared our marker lists to proteomic data by converting the list of transcripts to a list of encoding genes. We then compared these to the genes encoding proteins in the proteomic dataset. Any protein product of the encoding gene was considered a match. We used protein abundance data from 12-day-old flies (the second stable isotope labeling time point). Statistical significance was tested as above. In the case of Wan et al. [62], we compared a proteomic dataset directly to our head proteomics, but we also matched any protein product of a given gene rather than matching specific protein isoforms.

### Preparation of protein extracts for SDS-PAGE

#### RIPA buffer extraction

Ten to twelve heads from 1- or 10-day-old flies (equal numbers of males and females) were homogenized in 2x RIPA buffer with Sigma protease inhibitor cocktail (10 μL/head). RIPA buffer consisted of 50 mM Tris·HCl (pH 8), 150 mM NaCl, 0.5% NaDOC, 1% NP40, and 0.1% SDS. Samples were centrifuged for 5 min at 13,000 x *g*. An equal volume of 2x Laemmli buffer with β-mercaptoethanol (1:50) was added to each supernatant, and the supernatants were then boiled for 10 min and stored at −80°C.

#### Preparation of Triton-soluble and insoluble fractions

Samples consisted of 10 to 12 heads from 10-day-old flies or four whole flies (equal numbers of males and females). Samples were homogenized in 1x Triton lysis buffer (50 mM Tris-HCl [pH 7.4], 1% Triton X-100, 150 mM NaCl, 1 mM EDTA) with Sigma protease inhibitor cocktail (10 μL/head or 25 μL/fly). Homogenates were centrifuged at 15,000 x *g* for 20 min. The detergent-soluble supernatant was collected and mixed with an equal volume of 2x Laemmli buffer, and the same buffer was used to resuspend the detergent-insoluble pellet. All supernatant and pellet samples were boiled for 10 min. The detergent-insoluble protein extracts were centrifuged at 15,000 x *g* for 10 min, after which the detergent-insoluble supernatants were collected.

### Immunoblotting

Proteins were separated by SDS-PAGE on 4%-20% MOPS-acrylamide gels (GenScript Express Plus, M42012) and electrophoretically transferred onto PVDF membranes. Immunodetection was performed using the iBind Flex Western Device (Thermo Fisher, SLF2000). Antibodies were used at the following concentrations: 1:25,000 mouse anti-Actin (Chemicon/Bioscience Research Reagents, MAB1501), 1:500 mouse anti-ubiquitin (Santa Cruz, sc-8017), 1:2000 mouse anti-HA (BioLegend), 1:1000 mouse anti-mCherry (Invitrogen), and 1:500 mouse anti-GFP (BioLegend).

HRP secondary antibodies were used as follows: 1:500 to 1:1000 anti-mouse (BioRad, 170-6516), 1:500 to 1:1000 anti-rabbit (BioRad, 172-1019). Signal was detected using Pierce ECL Western Blotting Substrate (Thermo Scientific, 32106). Densitometry measurements were performed blind to genotype and condition using Fiji software [49]. Signal was normalized to Actin [114, 115]. For comparisons involving a genetic manipulation, the value for the control genotype was then set at 1. Normalized immunoblot data were log_2_-transformed to stabilize variance, and means were compared using Student *t* test. Significant results were defined as increases of at least 1.25-fold or decreases of at least 0.8-fold with *p* < 0.05. Each experiment was performed using at least three independent biological replicates.

### Fluorescent microscopy of *Drosophila* heads

One- or 10-day-old *Drosophila* (6-8 per condition, equal numbers of males and females) were anesthetized with CO_2_. Heads were removed and placed in Fluoromount aqueous mounting medium (Sigma F4680) on a glass slide with coverslip. Heads were then immediately visualized using a Leica MZFLIII microscope with Spot Insight Color Camera 3.2.0 and Spot Advanced software (exposure time 6-9 s). All cross-genotype comparisons were conducted on heads matched for gender.

### Drosophila behavior

All flies were housed in mixed-sex groups for at least 16 h to ensure mating.

#### Survival assay

Flies were initially housed in groups of 10 to 29, with 4-5 vials per gender per genotype. The number of flies in each vial (10-24) was recorded at 20 d of age, and flies were transferred to fresh food three times a week until they reached 40 d of age. Flies were again counted, and the percentage surviving to 40 d was calculated.

#### RING assay

Locomotor function was assessed using the Rapid Iterative Negative Geotaxis (RING) assay, modified from published protocols [71]. Flies were housed 8 to 15 per vial, separated by gender, and each genotype was represented by 6 to 12 vials. For the experiments involving expression of *Gba1b* in activated macrophages (*zfh1-GAL4 > UAS-Gba1b*), we used female flies, 13 d old; in the *GlcT* knockdown experiments, we used both male and female flies, 15 to 17 d old. Flies had a minimum of 24 h recovery time between carbon dioxide anesthesia and climbing. Six sets of flies were transferred to disposable plastic vials and loaded into a custom-built RING apparatus. Each set of six vials included both experimental and control vials. A Canon PowerShot ELPH 360 camera was used to record climbing height. The operator simultaneously allowed the RING apparatus to drop and pressed the camera button, which initiated a 3-second timer. Images were thus recorded ∼3 s after the flies were forced to the bottom of the vial. After a pause of 60 s, the procedure was repeated. At least five trials were performed for each set of vials. Climbing was scored using the first four trials without technical problems, by an experimenter blind to genotype. The position of each fly in the vial was manually marked on the image using Fiji software, and the average height climbed per vial was calculated for each trial.

#### Gut transit assay

Enteric nervous system function was tested using a procedure modified from Olsen and Feany [72]. Standard cornmeal-molasses food was dyed dark blue by adding two drops of blue food color (Safeway) per vial. The vials were then plugged and allowed to stand until dye was absorbed. Each genotype was represented by 4-5 vials. Flies were 15 d old in the *UAS-Gba1b* experiment and 20-22 d old in the *GlcT* knockdown experiment. Groups of 18-26 female flies were transferred to dyed food and allowed to feed overnight. The next day, a set of flies was quickly anesthetized with CO_2_, and the amount of blue dye visible through the ventral abdominal wall was scored for each fly under a dissecting microscope. This baseline measurement was performed to rule out confounding differences in food intake, and no such differences were observed. The flies were then transferred to regular fly food for 2.5 hours, after which blue dye in the abdomen was scored again. All scoring was done by an experimenter blind to genotype.

The assay was scored on a three-point scale (see Fig S7). No visible blue in the abdomen was scored 0, faint to moderate blue was scored 1, and intense blue was scored 2. Color intensity was scored by eye using an “area under the curve” approach, evaluating the total amount of dye visible in the abdomen. Flies with a score of 2 were considered to have impaired gut transit.

**Statistics**

Statistics were calculated using Microsoft Excel and GraphPad Prism. For ubiquitin-protein immunoblotting, in which only two conditions were compared, significance was calculated using Student’s *t* tests. As noted above, a criterion of 1.25-fold or 0.8-fold change in abundance was also applied to exclude biologically irrelevant differences.

Significance for RING and gut transit assays, in which four genotypes were compared, was calculated using one-way analysis of variance. If there were no significant differences in standard deviation, we used ordinary ANOVA with Tukey post-tests; if SDs differed, we used Brown-Forsythe and Welch ANOVA with Dunnett post-tests. Significance was set at *p* < 0.05 after correction for multiple testing.

Error bars represent standard error of the mean in all graphs. All experiments were performed using at least 3 biological replicates.

## Supplementary Figure Legends

**Figure S1. Schematic of analysis method comparing *Gba1b* mutant omics data to publicly available datasets.** The diagram illustrates our method of comparing *Gba1b* mutant RNA-Seq and proteomic data to available datasets on immune response and immune cell markers, showing two scenarios. See Materials and Methods for full details. The example is based on the marker list from the immune meta-analysis study [28], which includes 55 transcripts. (A) Chance-level overlap (no enrichment). Two transcripts appear both in the marker list and in the list of transcripts increased in abundance in *Gba1b* mutants. This is consistent with chance-level overlap, as 2/55 transcripts approximates the whole-dataset percentage of 3.5% transcripts with increased abundance. (B) Significant overlap. Twenty-eight transcripts appear both in the marker list and in the list of transcripts increased in abundance in *Gba1b* mutants. This is a statistically significant overlap, as 28/55 transcripts is 50.9%, more than 14 times the chance-level overlap of 3.5%.

**Figure S2. Interfering with expression of *Imd* pathway members does not ameliorate *Gba1b* insoluble ubiquitinated protein accumulation. (A)** Immunoblot of insoluble ubiquitinated proteins from heads of *Gba1b* mutants with and without *Relish* RNAi driven by *ppl-GAL4*. (B) Quantification of panel A for *Gba1b* revertant controls with vs. without *Relish* RNAi. The same comparison for *Gba1b* mutants is shown in Figure 2B. (C) Quantification of insoluble ubiquitinated proteins in heads from *Gba1b* mutants with and without loss of *PGRP-LC* function. (C) Quantification of insoluble ubiquitinated proteins from *Gba1b* mutants (whole flies) with and without the hypomorphic *imd^1^* mutation.

**Figure S3. Tests of macrophage markers and *GAL4* drivers for expression through 10 days of adult life.** Heads were matched for gender and imaged at 1 or 10 d of age, then evaluated for fluorescent signal in a distribution consistent with macrophages. Heads were imaged caudal side down, with the ventral aspect to the left. (A) Markers and GAL4 drivers that maintain macrophage expression through 10 d of adult life. (B) Well-known macrophage GAL4 drivers that lack visible macrophage expression at 10 d of age. (C) Other macrophage GAL4 drivers with expression that lack visible expression at 10 d of age.

**Figure S4. Confirmation of macrophage activation changes in 1- and 10-day-old *Gba1b* mutants using additional reagents.** (A) The *HmlΔ* promoter directly driving *dsRed* expression in control flies and *Gba1b* mutants gives the same result as *HmlΔ-GAL4*. (B) *VT17559-GAL4* contains promoter sequence from the *Lis-1* gene, an additional marker of macrophage activation. Like *zfh1-GAL4*, this marker is elevated in *Gba1b* mutants vs. controls at 10 d of age.

**Figure S5. Confirmation of rescue of macrophage activation using *HmlΔ-dsRed*.** The experiment was performed similarly to the rescue experiment shown in Figure 6A with *zfh1-GAL4* driving *UAS-Gba1b*. In this case, the *HmlΔ-dsRed* marker was substituted for *zfh1-lexA > tdTom*. Flies were 10 days old.

**Figure S6. Restoring *Gba1b* expression in activated macrophages rescues lifespan in *Gba1b* mutants.** *n* = 4-5 vials per group, 10-24 flies per vial at 20 d. \*\*\**p* < 0.005 by one-way ANOVA

**Figure S7. *Gba1b* mutants have abnormal enteric nervous system function.** (A) Illustration of gut transit assay adapted from Olsen and Feany [72]. Flies with a score of 2 (blue dye strongly visible in abdomen) were considered to have impaired gut transit. (B) Examples of control and *Gba1b* flies 2.5 h after switching from dyed to regular food. (C) Percent of flies retaining dye after 2.5 h. The *Gba1b* mutants and control flies used in panel C also bear *mCherry* RNAi. \*\*\**p* < 0.005 by *t* test.

**Figure S8. *Gba1b* expression driven in sLNv clock neurons by *pdf-GAL4* did not ameliorate macrophage activation in *Gba1b* mutants**. Macrophage activation was visualized using *zfh1-lexA* driving *lexAop-tdTom-HA* (*n* = 6-8 heads per genotype). Flies were 10 d old.

**Figure S9. Activated macrophage knockdown of *GlcT* using *zfh1-GAL4* caused impaired climbing in both control and *Gba1b* mutant flies**. Climbing was measured using the RING assay in male and female flies at 15-17 d of age. \*\*\**p* < 0.005 by one-way ANOVA.

